# Emergency Contraception Knowledge, Perceptions, and Barriers Among Married Saudi Men and Women: A Qualitative Study

**DOI:** 10.1101/2025.07.05.663316

**Authors:** Syed Irfan Karim, Farhana Irfan, Noura A. Abouammoh, Eiad Al Faris, Kamran Sattar, Abdullah MA Ahmed

**Affiliations:** Department of Family and Community Medicine, College of Medicine, King Saud University, Riyadh, Saudi Arabia; King Saud University Chair for Medical Education Research and Development, Department of Family and Community Medicine, College of Medicine, King Saud University, Riyadh, Saudi Arabia; Department of Medical Education, College of Medicine, King Saud University, Riyadh, Saudi Arabia

**Keywords:** Emergency Contraception, knowledge, barriers, qualitative research, male, female, Saudi Arabia

## Abstract

**Background:** Knowledge and use of contraceptives, including emergency contraception (EC), remain limited in Saudi Arabia due to cultural, religious, and gender-related influences. This qualitative study explores the knowledge, perceptions, and barriers related to contraceptive and EC use among married Saudi men and women.

**Methods:** Focus group discussions (FGDs) were conducted with purposively sampled married men and women at a university-affiliated government hospital in Riyadh, Saudi Arabia. Data was analyzed thematically to explore participants’ beliefs, attitudes, and experiences regarding contraceptive and emergency contraception use.

**Results:** Men expressed positive attitudes toward EC use. Overall, understanding of contraception was limited. The identified barriers for use of EC were lack of information, misconceptions, fear of side effects, social norms, and religious beliefs. Participants highlighted the importance of enhancing education through premarital counseling and public awareness programs.

**Conclusions:** This study highlights the need for improved education on contraception. Given the influence of male perspectives on contraceptive use, increasing awareness by culturally appropriate, gender-sensitive educational strategies to dispel myths and promote informed contraceptive decision-making could be a valuable intervention to enhance contraceptive uptake in Saudi Arabia

## Introduction

Globally, approximately 40% of all pregnancies are unintended [1,2], with approximately 30% attributed to contraceptive failure [3]. Many couples desire for smaller families often face the challenges in preventing unintended pregnancies. It could be either due to method or user failure. Emergency contraception (EC) is an option to prevent pregnancy after unprotected or inadequately protected intercourse [4]. Despite its documented benefits, its usage continues to be lower in the Muslim countries compared to the West [5,6].

It’s essential to note that views on contraception, including emergency contraception, can vary among individuals within any religious community, and opinions may be influenced by cultural, regional, or individual differences. Bridging this gap requires addressing factors influencing awareness, access, and cultural perceptions surrounding emergency contraception in Muslim communities.

In the Middle East and North Africa region (MENA) (especially the Arab world) the awareness and use of EC is markedly lower than in most Muslim countries and the West [7,8]. The preferences for contraceptive methods in this region are also different than those in the West [9]. Effective contraception is crucial in the Muslim world due to religious and cultural prohibitions against pregnancy termination [10–12]. EC use is a prominent topic of debate, and controversies exist that may be due to lack of knowledge and conflicting views about contraception and abortion [13]. Additionally, contraception is often considered primarily a women’s issue [14]. Contraceptive research usually examines women’s perspective on EC, it is equally important to acknowledge vital role men play in preventing unintended pregnancies [15].

Partners jointly decide upon the desired family size. However, husbands often play a crucial role in global fertility decision-making [16]. A husband’s involvement in family planning positively affects contraceptive use. Individuals’ sexual behavior and preferences are influenced by factors such as personal attitudes, beliefs, community gender norms, context, partners influence over method selection and EC conceptualization [17,18]. Despite widespread contraceptive utilization, barriers such as method, side effects, lack of information, cost and cultural factors hinders consistent use [19], which can lead to periods of unprotected intercourse and unintended pregnancy.

Most studies on EC, globally and particularly in Arab region, adopt a quantitative design, which limit in-depth probing and understanding of the phenomenon. A recent qualitative study has explored Muslim women’s awareness and experiences with family planning in Saudi Arabia [20]. However, there is a gap for specific inquiry regarding EC; particularly involving men. Literature has shown that the current knowledge of EC among men is limited in the region [21]. Hence, their attitudes, behaviors, and awareness is crucial for understanding and promoting contraceptive use. An understanding why individuals avoid EC is critical to address unmet needs and promote contraceptive use.

Based on the research gaps discussed above, this study was designed to explore knowledge, attitude and barriers about EC. In this study the Health Belief Model (HBM) along with conceptual framework of Hall was utilized.

The constructs of the model offer a framework for evaluating patients’ behavior patterns and needs within complex social, environmental and reproductive contexts. It considers humans as logical thinkers with multidimensional approach to decision making and health behavior choices [22,23].

### Rationale for the study

Theory-driven strategies to prevent unintended pregnancy are currently needed [24]. Limited qualitative research in the region exists on the perceptions of both men and women regarding EC. More specifically, understanding their thoughts and needs on contraceptive use is important to improve reproductive health outcomes. To address the research gaps, this study explored the knowledge, perceptions, modifying and enabling factors (barriers and concerns) related to EC among Saudi men and women.

## Methodology

### Study setting

This study was conducted in King Saud University Medical City (KSUMC), in the capital city Riyadh, Saudi Arabia.

### Study design

This was a qualitative phenomenological study eliciting knowledge, views, and experiences of men and women about emergency contraception (EC) using focus group discussions (FGDs).

### Target population

The population targeted was Saudi men and women attending primary healthcare clinics in King Saud University Medical City (KSUMC).

### Sampling and recruitment

A convenient sampling was used to select participants based on the eligibility criteria. The eligibility criteria included being a male or female within the reproductive age group and consenting to participate in the study. A sample of 6-8 patients from both sexes was selected for distinct focus group discussions.

Recruitment took place at the primary care clinics (PCCs) of KSUMC in Riyadh.

Patients were identified through the consultation patient lists, and the research team approached them by phone two days prior to their scheduled visit. A detailed explanation of the study was provided. After obtaining written consent, participants were contacted on the scheduled appointment day, either just before or after their consultation. Upon reporting to the reception desk, a research team member reconfirmed their willingness to participate and sought written consent. An information sheet explaining the aims of the study was distributed to the patients in the waiting area by the research team members. Furthermore, researchers meticulously explained the study’s methodology, addressing individuals’ consideration’s such as physical fitness or time constraints.

### Data collection

The data needed to achieve the objectives of this study included knowledge, perceptions, modifying factors, and enabling factors (such as barriers and concerns) related to EC among Saudi men and women. While various methods could have been used to collect this data, focus group discussions were chosen for two primary reasons: they provide a comfortable environment for participants to share their perspectives alongside others of the same gender, reducing power imbalances between researchers and participants, and they also encourage participants to recall and reflect on their views more effectively. The patients were informed about the aims of the study, anonymity and confidentiality was assured. The FGDs were conducted in the nearby seminar room of the PCCs. The length of the interviews was decided to be kept for 60 minutes. Data collection was continued until reaching data saturation. The FGDs lasted for an average duration of 60-90 minutes.

All FGDs were conducted in person by the researchers. (NA) in the presence of another member of the research team (FI) and (EA) along with (SI), conducted the focus group discussions for female and male groups separately. Researchers were faculty in the College of Medicine. NA and EA has special interest in qualitative research, and extensive experience in qualitative methodologies, data collection, and analysis techniques. Keeping in mind the sensitivity of the topic and the culture, female researchers conducted the interviews for women and the male researchers for men. Participants were not known to the interviewers before the data collection sessions. Only the participants and researchers were present during the focus group discussions. The completion of demographic information preceded all the interviews. Refreshments were provided during the focus group discussions to for a relaxed and informal atmosphere for the participants. At the beginning of the interview, the researchers written consent for voice recording was obtained. Before starting the discussion, it was made clear to the patients that their withdrawal from the study was allowed at any time without any implications on the health service received. A semi-structured FGD topic guide designed by the was used during the interviews. It was formulated in English and translated into Arabic by a native Arabic researcher using forward and back translations. The discussions were conducted in Arabic, as it was patient’s preference. Additionally, participants were encouraged to respond openly and share their opinions and experiences. However, they were not pressured to answer questions or share sensitive information.

Data collection was continued until data saturation. Conversations were audio-recorded with the participants’ consent. It was explained to the participants that they have the right to ask the researcher to stop the recording before the focus group discussion or at any time during its progress. Four (4) FGDs were conducted, as recommended by a systematic review to achieve data saturation [25.] Saturation was achieved when participants’ responses during the interviews no longer revealed new information.

### Topic guide

The Focus Group Discussion (FGD) topic guide was developed based on findings of previous studies and amended according to the culture and covered the area of knowledge, perception, and utilization of emergency contraception (EC), as well as motivators and barriers. Open ended interview questions were designed that allowed participants to express their perspectives freely, reducing the researcher’s influence on their responses.

The FGD topic guide included questions exploring experiences with modern and traditional contraceptives, reasons for nonuse, and the factors influencing contraception decision making. The interview guide incorporated the questions shown in the Table :1

**Table 1:**
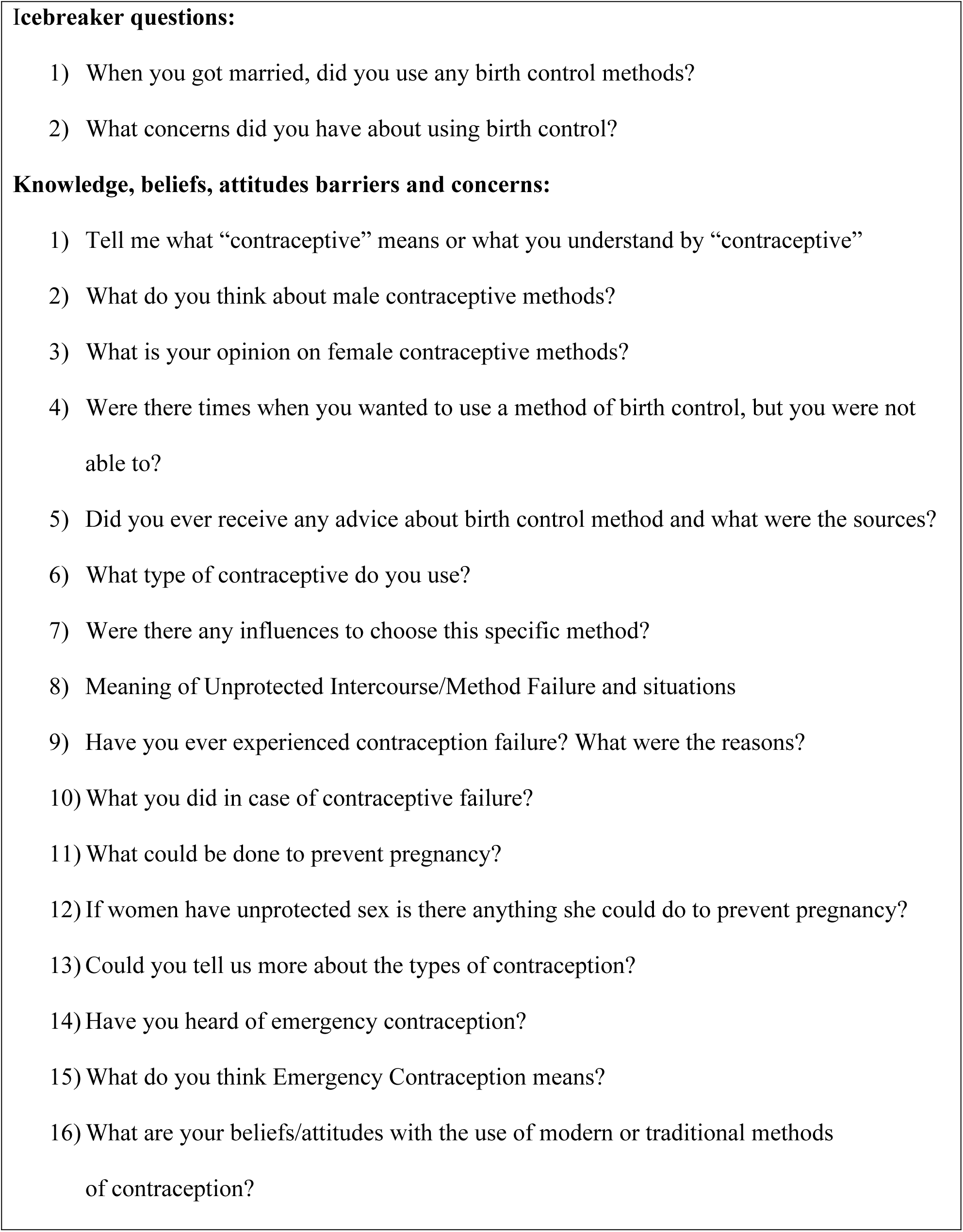
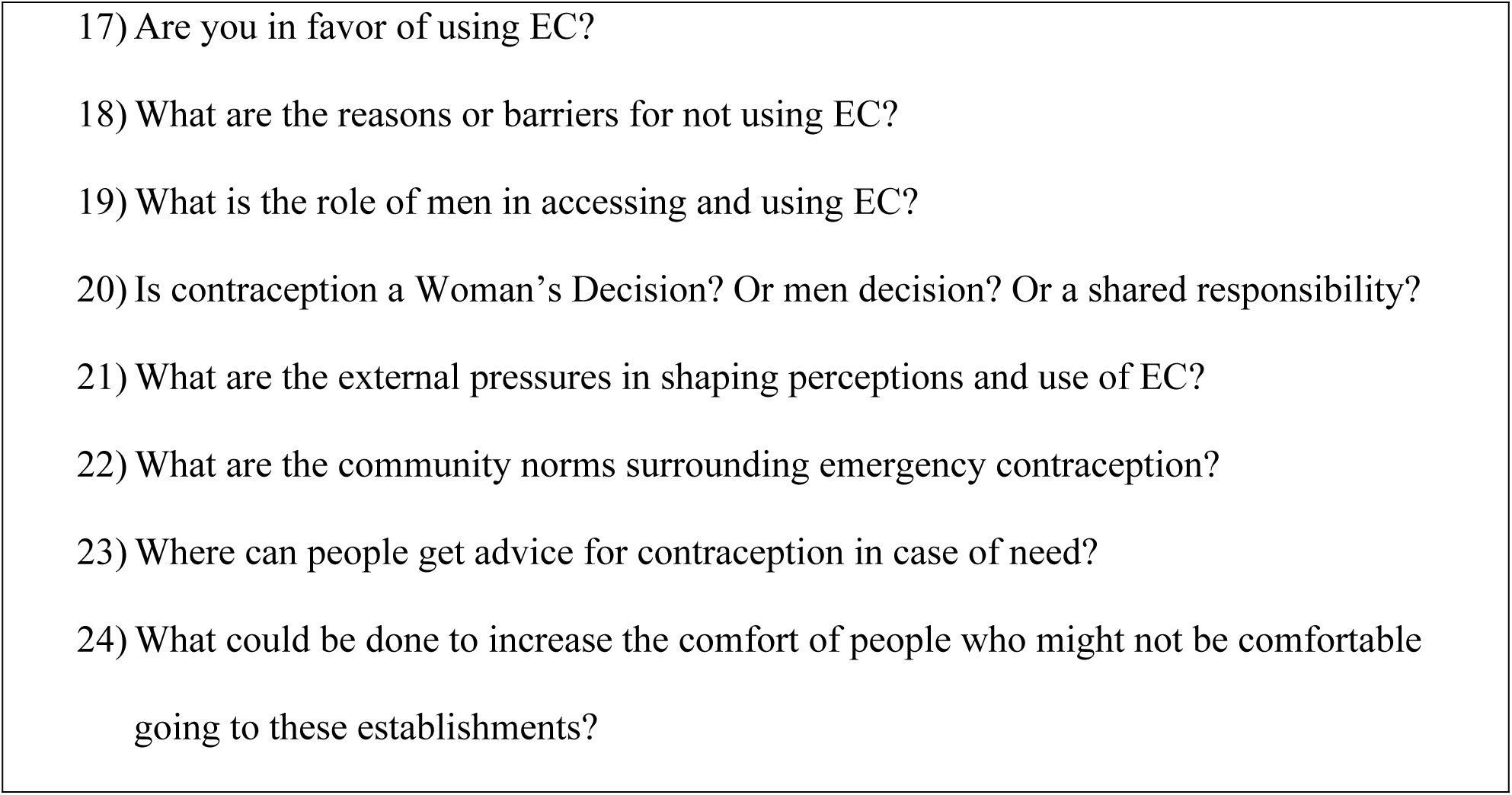
List of questions used during FGDs.

### Data analysis

We used the thematic analysis method suggested by Ritchie and Spencer [26] which offers a structured, step-by-step framework for analyzing data. The following outlines the process involved during our thematic analysis.

#### 1 -Familiarization

Listening to recordings; transcribing verbatim; translating and reading and re-reading through all data to gain a holistic overview of the dataset and be familiar with the range, depth and diversity of the collected data was done by NA.

#### 2 -Descriptive coding

The computer-assisted qualitative data analysis software NVivo 10 was used for transcription and coding. The research team read the same transcripts and agreed on an initial coding frame, which was applied to other transcripts, with the flexibility to make it possible to add other codes.

#### 3 -Basic analysis

At this stage, themes and subthemes emerged as codes were logically grouped to address the research question. Data were removed from their original context and reorganized according to the relevant thematic orientation.

#### 4 -Interpretation

This was the final stage where the key themes were developed and quotations used to provide supporting evidence. It involved exploration of the thematic network in a systematic way. It encompassed mapping the range and nature of phenomena, fully exploring each theme and finding associations between themes. Data collection and analysis continued until theoretical saturation. To enhance the robustness of the complete process, we utilized the Consolidated Criteria for Reporting Qualitative Studies (COREQ) checklist. By utilizing COREQ, we were able to systematically address potential biases, improve the credibility of our findings, and facilitate the reproducibility of our research. The completed checklist is available in **S 1 Appendix**.

### Ethical considerations

The study was conducted based on the Helsinki Declaration and was approved by the Institutional Review Board (IRB) of KKUH, King Saud University (Project No. E-21-5719).

There was no relationship between the researchers and the participants. Written informed consent of individual participants was taken and were assured of the confidentiality of their data.

## Results

### Participants Demographics

#### Analysis of the data

A total of_Sixteen participants were recruited four focus group discussions, until data saturation. Most of the participants were in the age groups of 30-40 (50%), and all were married. IUD’s were the most commonly used modern contraceptive methods (50%), followed by OCPs (25%); whereas periodic abstinence was least used method of contraception (6%). **Table 2** provides an overview of the demographic details of the participants.

**Table 2:** Demographic details of the participants.

| Participants Male (M) | Age | Currently married | Contraceptive use | Occupation |
| --- | --- | --- | --- | --- |
| M1 | 36 | Yes | Patches | Office Job |
| M2 | 41 | Yes | IUD | Manager |
| M3 | 36 | Yes | IUD | Secretary |
| M4 | 35 | Yes | OCP | Secretary |
| M5 | 48 | Yes | IUD | Office Job |
| M6 | 45 | Yes | IUD | Manager |
| M7 | 43 | Yes | OCP | Office Job |
| M8 | 38 | Yes | NIL | Secretary |
| Participants Female (F) | Age | Currently married | Contraceptive use | Occupation |
| F1 | 29 | yes | None ( don't have kids) | House wife |
| F2 | 27 | yes | OCP | Nurse |
| F3 | 31 | yes | IUD | Nurse |
| F4 | 33 | yes | Withdrawal | Nurse |
| F5 | 28 | yes | IUD | Nurse |
| F6 | 41 | yes | IUD | Nurse |
| F7 | 36 | yes | OCP | Security Job |
| F8 | 39 | yes | IUD | Security Job |

### Themes and subthemes

Two main themes were identified: Knowledge gaps and misconception about contraception and perceived barriers about contraception. (Table :3)

**Table 3 :** Themes and subthemes.

| Themes | Subthemes |
| --- | --- |
| Knowledge Gaps and Misconceptions | 1. Limited awareness<br>2. Confusion and misconception. |
| Barriers about Contraception. | 1. Perceived Religious prohibitions and ethical concerns<br>2. Moral dilemma's<br>3. Stigma and social Judgement<br>4. Gender dynamics and decision making power |

### Knowledge Gaps and Misconceptions

The study began with exploring the participants’ knowledge of contraception. Participants knowledge about contraceptive methods was primarily for pills (OCP’s). All participants were aware of condoms as a male and OCP’s as female contraceptive method; however, other methods were not widely known. Only 3 women knew about hormonal injections for contraception. Few mentioned other methods and were able to name some commonly available options. While EC as an option for contraception and the appropriate time frame for it was known by only one female participant. She stated that it is available in Saudi pharmacies with a prescription. Two other female participants had heard of EC but lacked sufficient knowledge about it.

Few men had idea of other methods.

> *“The pills, the patches, intrauterine device”* **M3**

> *“There is a thing that they put under the skin and also condoms or refrain from intercourse in certain time of the cycle”* **M4**

> *“Vaginal douche after intercourse can help”* **F1**

> *“There are some pills men could use…I read about it but I do not think it is available in Saudi Arabia”* **M4**

> *“There are some pills and injection men could use as contraceptives…I don’t have enough information about, except that they are hormonal, and may cause mood swings”* **F4**

> *“It is a pill that the women take if she is doubting pregnancy, that’s all I know”* **F2**

> *“There is some operation men can undergo that is equivalent to tubal ligation in women!”* **F4**

However, none of the male participants had heard about EC. However, when the idea of emergency contraceptive was explained to them, they supported its use for many reasons:

> *“It may be looked at as a savior if any contraception failed.”* **M3**

> *“It is good for family planning and also for delaying pregnancy due to financial and accommodation issues”* **M4**

> *“you need to space between children to be able to give the love and attention each child deserve”* F4

Misconceptions and myths about contraceptives were prevalent among women, often influenced by stories shared within their communities. For instance, some believed that hormonal contraceptives could lead to infertility.

Some male participants also supported this view,

> *“The women may not be able to get pregnant in the future if she previously used contraception”* M2

Most participants preferred to avoid hormonal contraception until after having their first child. until later after having a child.

> *“I got married 10 years ago, we only started using contraceptives after we had our son”* **M2**

It was also recommended by one of the participants to change the type of contraception women use to prevent infertility because of contraception use. She noted:

> *“You have to change the hormone or stop using them every 3 to 4 months because the body gets dependent on the hormone and react differently to it in the system and then this may cause infertility”* **F6**

> *“For Newly married couples it may affect the wife’s fertility. …my friend used contraception during the initial six months of her marriage. Then she was infertile for a period of 5 years”* **F1**

When asked about the perceived reason for this phenomenon, she replied:

> *“using hormonal contraceptives before getting pregnant for the first time causes weakness of ovulation”* **F1**

Another one added:

> *“When new hormones are introduced into your body, all the natural hormones get disturbed and this may cause temporary or permanent infertility”* **F6**

A few female participants preferred using herbal remedies as an alternative to emergency contraception (EC).

> *“I prefer taking natural products but not chemicals. I have an experience with a herb called (kaf Mariam/Mary’s palm) in the delivery of my second baby and I can tell it contracts the uterus effectively”* **F7**

Participants did not think of the copper IUD as EC because they viewed it as “planned,” preventative method, not fitting within their definition of EC.

> *“I never thought of that Intrauterine Device (IUD) was always a routine contraceptive. I didn’t know that it was an emergency Contraceptive method as well.”* **F 8**

> *“Copper intrauterine device is the best! Why would a women introduce her body to more hormones!?”* **F6**

> *The perception for contraception showed mixed feelings; men considered it mainly for financial reasons*.

> *“It is good for family planning and also for delaying pregnancy due to financial and accommodation issues” M6*

> *“for newly married couples, pregnancy is ok; but the situation may not be the same for those who already have six children, so it can vary depending on individual circumstances”.* **M 2**

> *“I don’t think contraception is good”.* **M4**

> *“It’s good if the wife has medical conditions preventing her from taking OCPs”.* **M4**

> *“A temporary solution to control pregnancy”.* **M3**

> *“Preventing pregnancy for reasons like raising children and breastfeeding”.* **M3**

Men primarily relied on their spouses as their main source of information on contraception, viewing their wives as responsible for contraceptive knowledge.

> *“I think the wife is the one who’s responsible for getting the information”.* **M 4**

> *“First my wife asks expert people like her mother and older sisters, If she doesn’t find an answer she then seeks medical advice”.***M4**

Women did not trust internet as a source of information to choose the best type of contraceptive method for them.

> *“Women’s stories on the internet are unreliable, as each woman responds differently to different contraception methods. Even sisters may have varying experiences with different methods”* F5

Only a few women preferred going to a healthcare provider for contraceptive advice.

Few females however, visited the **doctor** before their marriage to consult them about their option of contraception

However, most male participants use the **internet** to educate themselves about types of contraception. Some other participants ask the **pharmacists** about different kinds of contraception that can be used and share their information with their wives.

The participants’ narratives highlights’ an overall positive attitude towards the use of emergency contraceptive pills. This perspective was reflected by the following statements:

> ***“*** *I strongly support it, as someone who has a disabled child, I understand the importance of family planning. It’s crucial to think about the consequences before getting carried away in the moment”.***M5**

> *“It is a good solution for those who cannot deal with having a new baby or who cannot pay its expenses”* **F2**

> *“It may be looked at as a savior if any contraception failed”.* **F4**

> *“If I knew about EC 2 years ago I would have used it. I really did not want to get pregnant at that time”* **F3**

### Beliefs and Barriers

#### Gender dynamics and decision making power

Male dominance in decision making:

Men often have limited knowledge but significant influence on contraceptive decisions. Although, the contraception was regarded a women affair by men; at the same time, they seemed to dictate women on family size and contraceptive use.

> *“Definitely, the husband has a big role in the decision whether to have a baby or not”.* **F 6**

#### Women’s Autonomy

Women face challenges in discussing EC due to male partners’ opposition. Women also mentioned that discussing the use of emergency contraception (EC) with their partner was unnecessary, as having a child would have a greater impact on the mother’s life than on the father’s.

> *“He may say: it is a gift from Allah (God), but we are the one who takes care of the baby’s life”* **F4**

#### Male partner disapproval of male contraceptives

Some women expressed frustration about being the ones expected to take responsibility for contraception. When they asked their husbands if they would consider using male contraceptive methods, they were met with refusal.

> *“Men do not agree to use contraception, there is no way my husband would agree to take part in using even condom”* **F3**

Another participant added:

> *“Men may think that if they used contraceptives, their masculinity may get affected. This is true for all eastern men…it is the culture”* F4

#### Stigma and Social Judgment

Societal expectations and fear of negative perceptions from the community discourage open discussions about sexual issues and contraception, leading to reluctance to seek help.

> *“It’s not commonly understood in the community”.* **M2**

> *“You may get criticized by your relatives”.* **M3**

A small number of male participants found it difficult to use condoms for cultural related issues, for example, one participant noted:

> *“I do not use condoms. It is kind of embarrassing in front of my wife. People in the community are not used to using it”* **M4**

Another one added:

> *“The intimacy is effected when using condoms”* **M5**

In line with this, most female participants preferred their partners’ comfort, preference and convenience. They felt that methods like condoms, withdrawal and abstinence were not ideal as it affects their partners’ pleasure. One mentioned:

> *“We use the withdrawal method. But my husband has to put his mind in not forgetting to withdraw at the right time, which may affect the whole experience”* **F4**

While all participants agreed on the importance of discussing contraceptive use with their partners, all male participants reported that they leave the choice of contraceptive method to their wives. In line with this, female participants felt they knew what works best for their bodies and therefore informed their husbands of their chosen method, rather than engaging in mutual discussion.

#### Side effects

The reluctance to use modern contraceptives was attributed to side effects. Male participants noted that hormonal OCP may cause mood swings.

> *“Pills may change the women’s’ mood. That’s why my wife used skin patches”* **M1**

> *“My wife started taking OCPs for 9 years, but she stopped now after she suffered from psychological side effects”.* **M5**

> *We couldn’t continue on the OCPs due to mood changes.* **M8**

> *“Ectopic pregnancy can happen with the IUD use, or even a complete pregnancy”* **F 4**

> *Few participants were hesitant to use EC because of the fear of fetal deformities. “What if it affected the baby and caused deformities without causing abortion!?”* **F8**

> *“Some people wouldn’t use it thinking it may cause fetal malformation if it fails”.* **F7**

Others, however, believed that contraception could cause infertility and preferred not to risk using it until after having at least few children.

> *“I got Married in 2007, and didn’t use contraception for the first 10 years of marriage but after 4 kids my wife started taking pills.” **M*** **7**

> *“It may cause infertility”.* **F 3**

Effect on babies like fetal deformities.

> *“It (OCP) may cause birth defects”* **F7**

While using contraceptive, all participants will accept their pregnancy if it happens and would not take any procedure to terminate their pregnancy.

#### Religious concern

Most female participants did not like the idea of the EC and considered using it as intended abortion. One mentioned:

> *“It is Haram (religiously forbidden), I would not use it. It is killing a soul”* **F 2**

Some female participants mentioned that they might wait for a Fatwa (a religious ruling) on the use of emergency contraception (EC), believing that religious scholars would approve it due to the high-cost living conditions.

#### Role of Education and Awareness Campaigns

Participants thought that efforts should be done to educate the community about this kind of contraception via:

> *“SMS messages from the MOH, through pharmacists…”* **M1**

> *“They should integrate contraceptive education into premarital clinics—husbands need to be informed.” –* **F4**

> *It’s a new thing to the community but with time they will accept it*. **M 6**

Participants suggested increasing the awareness about EC among people through the OBGYN clinics and social media to prevent unwanted pregnancies.

Other considerations mentioned by female participants to encourage the uptake of EC were:

> *“What is the rate of success of these pills and if it is Haram”* **F2**

Female participants expressed a strong need for education on the various types of contraception and what to expect from their use. All participants recommended the establishment of a premarital or family planning clinic specifically for men, where they could also receive education about contraceptive options. Additionally, two participants emphasized the importance of informed decision making.

> *“Nowadays, women go straight to the pharmacy and choose a contraceptive method… but they need to be educated about their options.” –* **F2**

## DISCUSSION

This study identified significant gaps in contraceptive knowledge, misconceptions, and social factors that shape contraceptive behaviors in Saudi Arabia, particularly among married men and women.

A consistent theme that emerged from the focus group discussions was the limited knowledge/awareness of contraceptive options. Notably, men exhibited lower levels of awareness about contraceptive options compared to women, with emergency contraception (EC) being especially poorly understood.

Only a handful of participants had heard of EC, and only one was able to articulate its appropriate use. Amon men none had prior knowledge of EC, this is consistent with findings from other studies where low awareness to contraceptive methods results in poor contraceptive intake [26].Interestingly, once the concept was explained, both men and women expressed supportive attitudes, viewing EC as a helpful option for family planning and in the case of contraceptive failure. This finding suggests that educational interventions may significantly improve acceptance and use of EC, provided they are culturally sensitive and address misconceptions.

Misconceptions about hormonal contraceptives, including fears of infertility and fetal deformities, mood swings and psychological distress were common among participants and were often influenced by anecdotal tales circulating in their communities. Participants, especially women believed that use of hormonal contraception could lead to infertility. This fear can be a key hindrance to contraceptive use in the region [20]. Similar concerns were raised by several studies conducted elsewhere about the emergency contraceptive side effects [27–29]. Reliance on sources of information, such as the internet, pharmacists, or family members, rather than healthcare providers, may have contributed to misconceptions and inadequate knowledge, potentially acting as a barrier to effective contraceptive use. Gender dynamics emerged as a critical factor influencing contraceptive decision-making. Men often viewed contraception as a responsibility for women but still played a significant role in family planning decisions, reflecting cultural norms that emphasize male authority. Women shared their frustration about their male partners refusing to use male contraceptive methods. Participants also reported stigma and embarrassment associated with male contraceptive methods such as condoms. Men feared that using such methods might impact their masculinity or social standing. This finding is particularly important as it highlights a cultural resistance to male participation in family planning, further placing the burden of contraception on women.

Similar perception among married couples have been reported in studies from other regions, where pregnancy prevention is regarded as a woman’s responsibility [30–32]. This limitation on women’s autonomy in reproductive decision-making is consistent with findings from other research within the region [33,34].

A possible reason could be women seek care from female health providers, which limits male participation and discussion around family planning. This is further compounded by cultural and social norms. It is important to understand men’s perceptions and attitudes toward contraception and to include men in family planning services. By providing male-centered care, men can feel more comfortable discussing reproductive health with male healthcare providers, addressing their needs at the community level. Additionally, religious beliefs played a pivotal role in shaping attitudes toward EC, with some female participants equating its use to abortion and considering it “haram” (forbidden). Some female participants might use it if there is a fatwa (a religious ruling) by religious authorities. Other studies have similarly noted that Saudi women perceive contraception as conflicting with Islamic beliefs and principles [35,36]. This reflects the importance of religious influence in making reproductive choices, suggesting that collaboration with religious scholars could strengthen the cultural acceptance and effectiveness of contraceptive education programs. To address these cultural and religious concerns, educational interventions must be culturally sensitive and utilize mass media to correct myths and shift social norms [35].

Additionally, community mobilization and personalized counseling where satisfied contraceptive users share their experiences, are also effective strategies [37].

Lastly, study findings emphasize that participants recommended premarital counseling and community based education, particularly for men, to improve contraceptive knowledge. Establishing family planning clinics specifically for men and adding contraceptive education into premarital counseling programs could also promote shared decision-making.

These recommendations align with existing literature, which highlights the role of comprehensive educational initiatives in improving awareness and empowering individuals to make informed choices and use contraception more effectively [38–39].

## Conclusion

This study contributes to the existing literature on contraception knowledge and attitudes by highlighting the limited knowledge especially among men and their tendency to rely solely on their partners for contraception and its issues. The findings highlight the important need for educational programs that include men, as their awareness and involvement are essential for the successful use of contraception. Future research should focus on bridging the gaps to ensure comprehensive reproductive health care for all.

### Strengths

To the best of our knowledge, this is the first time that perceptions, attitudes, misconceptions, knowledge and barriers about EC among married men and women in Saudi Arabia has been reported. This study was conducted in one of the prominent hospitals in the capital city of the kingdom of Saudi Arabia. A key strength of this study is the inclusion of both Muslim men and women. Previous research has primarily focused on the views of Muslim women regarding contraception and unintended pregnancy.

### Limitations

This study has a number of limitations that should be noted. Because our study population comes from a specific geographic area and represents average income women and men, our findings are not generalizable to the experience of Saudi Arabian women /men from more diverse sociodemographic backgrounds. Also, given that our primary recruitment was through university-based hospital clinics, our study population may represent women and men who are generally more comfortable accessing health care services in general.

Another possible limitation of the study is that interviews were conducted by two different interviewers, which may have introduced variability in data collection and influenced the consistency of responses. This study was conducted in only one of the prominent hospitals in the capital city of Saudi Arabia, covering a small group not representative for all men and women in urban setting. Participants from other cities in Saudi Arabia may have more conservative views regarding contraception. Therefore, generalizing the results of this study to other settings must be done with caution.

## Abbreviations

EC: Emergency contraception.
FGD: Focus Group Discussion.
HBM: Health Belief Model.
PCC: Primary care Clinics.
COREQ Checklist: Consolidated Criteria for Reporting Qualitative Studies.

## Consent for publication

Not applicable

## Competing Interests

The authors declare that they have no competing interests.

## Funding

The author extends their participation to the Deanship of Scientific Research, King Saud University for funding through Vice Deanship of Scientific Research Chairs, Research Chair of Medical Education and Development.

## Authors’ contributions

**Conceptualization:** Syed Irfan Karim, Farhana Irfan

**Data curation:** Farhana Irfan, Syed Irfan Karim, Noura A. Abouammoh, Eiad Al Faris Kamran Sattar, Abdullah MA Ahmed.

**Formal analysis:** Syed Irfan Karim, Farhana Irfan, Noura A. Abouammoh, Eiad Al Faris Kamran Sattar, Abdullah MA Ahmed.

**Funding acquisition:** Syed Irfan Karim, Abdullah MA Ahmed,

**Investigation:** Syed Irfan Karim, Farhana Irfan, Noura A. Abouammoh, Eiad Al Faris

**Methodology:** Syed Irfan Karim, Farhana Irfan, Kamran Sattar.

**Project administration:** Syed Irfan Karim, Farhana Irfan

**Resources:** Syed Irfan Karim, Farhana Irfan, Noura A. Abouammoh, Eiad Al Faris.

**Software:** Noura A. Abouammoh, Abdullah MA Ahmed

**Supervision:** Syed Irfan Karim

**Validation:** Syed Irfan Karim, Farhana Irfan

**Visualization:** Syed Irfan Karim, Farhana Irfan,

**Writing** – **original draft:** Syed Irfan Karim

**Writing** –**review & editing:** Syed Irfan Karim, Farhana Irfan, Noura A. Abouammoh

## Supporting information

**S 1 Appendix**

## Notes

### Competing Interest Statement

The authors have declared no competing interest.

## References

1. S.K. Kathpalia. Emergency contraception: Knowledge and practice among women and the spouses seeking termination of pregnancy. Med J Armed Forces India. 2016;72 (2):116-9. available from: https://www.ncbi.nlm.nih.gov/pubmed/27274610

2. Yen S, Parmar DD, Lin EL, Ammerman S. Emergency Contraception Pill Awareness and Knowledge in Uninsured Adolescents: High Rates of Misconceptions Concerning Indications for Use, Side Effects, and Access. J Pediatr Adolesc Gynecol. 2015 Oct;28(5):337-42. doi: 10.1016/j.jpag.2014.09.018. Epub 2014 Oct 8. PMID: 26148784.

3. Bearak J, Popinchalk A, Alkema L, Sedgh G. Global, regional, and subregional trends in unintended pregnancy and its outcomes from 1990 to 2014: estimates from a Bayesian hierarchical model. Lancet Glob Health. 2018 Apr;6(4): e380-e389. doi: 10.1016/S2214-109X(18)30029-9. Epub 2018 Mar 5.

4. Emergency contraception. Practice Bulletin No. 152. American College of Obstetricians and Gynecologists. Obstet Gynecol 2015;126:e1–11. Available from: https://www.acog.org/clinical/clinical-guidance/practice-bulletin/articles/2015/09/emergency-contraception

5. Emergency contraception: key facts. World health organization, 2018. Available from: https://www.who.int/news-room/fact-sheets/detail/emergency-contraception

6. Sweileh WM, Zyoud SH, Al-Jabi SW, Sawalha AF. Worldwide research productivity in emergency contraception: a bibliometric analysis. Fertil Res Pract. 2015 May 5;1:6. doi: 10.1186/2054-7099-1-6. PMID: 28620511; PMCID: PMC5415191.

7. Karim SI, Irfan F, Rowais NA, Zahrani BA, Qureshi R, Qadrah BH. Emergency contraception: Awareness, attitudes and barriers of Saudi Arabian Women. Pak J Med Sci. 2015 Nov-Dec;31(6):1500-5. doi: 10.12669/pjms.316.8127. PMID: 26870124; PMCID: PMC4744309.

8. Marafie N, Ball DE, Abahussain E: Awareness of hormonal emergency contraception among married women in a Kuwaiti family social network. Eur J Obstet Gynecol Reprod Biol 2007, 130: 216–222. 10.1016/j.ejogrb.2006.05.023.

9. Sedgh G, Singh S, Hussain R. Intended and unintended pregnancies worldwide in 2012 and recent trends. Stud Fam Plann. 2014 Sep;45(3):301-14. doi: 10.1111/j.1728-4465.2014.00393.x. PMID: 25207494; PMCID: PMC4727534.

10. Shapiro GK. Abortion law in Muslim-majority countries: an overview of the Islamic discourse with policy implications. Health Policy Plan. 2014 Jul;29(4):483-94. doi: 10.1093/heapol/czt040. PMID: 23749735.

11. El Hamri N. Approaches to family planning in Muslim communities. J Fam Plann Reprod Health Care. 2010 Jan;36(1):27-31. doi: 10.1783/147118910790291019. PMID: 20067669.

12. Al-Matary, A., Ali, J. Controversies and considerations regarding the termination of pregnancy for Foetal Anomalies in Islam. BMC Med Ethics 15, 10 (2014). 10.1186/1472-6939-15-10.

13. Al-Matary, A., Ali, J. Controversies and considerations regarding the termination of pregnancy for Foetal Anomalies in Islam. BMC Med Ethics 15, 10 (2014). 10.1186/1472-6939-15-10

14. Chng CL. The male role in contraception: implications for health education. J Sch Health. 1983 Mar;53(3):197-201. doi: 10.1111/j.1746-1561.1983.tb07820.x. PMID: 6552331.

15. Marcell AV, Waks AB, Rutkow L, McKenna R, Rompalo A, Hogan MT. What do we know about males and emergency contraception? A synthesis of the literature. Perspect Sex Reprod Health. 2012 Sep;44(3):184-93. doi: 10.1363/4418412. Epub 2012 Jul 12. PMID: 22958663.

16. Tilahun T, Coene G, Luchters S, Kassahun W, Leye E, Temmerman M, et al. (2013) Family Planning Knowledge, Attitude and Practice among Married Couples in Jimma Zone, Ethiopia. PLoS ONE 8(4): e61335. 10.1371/journal.pone.0061335.

17. Habibzadekh F. Contraception in the Middle East The Lancet 2012; vol 380. Available from: https://www.thelancet.com/pb/assets/raw/Lancet//pdfs/Sep12_MiddleEastEd.pdf

18. Schölmerich VL, Kawachi I. Translating the Social-Ecological Perspective Into Multilevel Interventions for Family Planning: How Far Are We? Health Educ Behav. 2016;43(3):246-255. 10.1177/1090198116629442

19. Wright RL, Fawson PR, Frost CJ, Turok DK. U.S. Men’s Perceptions and Experiences of Emergency Contraceptives. Am J Mens Health. 2017 May;11(3):469-478. doi: 10.1177/1557988315595857. Epub 2015 Jul 17. PMID: 26186949; PMCID: PMC5675232.

20. Alomair N, Alageel S, Davies N, Bailey JV. Muslim women’s views and experiences of family planning in Saudi Arabia: a qualitative study. BMC Womens Health. 2023 Nov 25;23(1):625. doi: 10.1186/s12905-023-02786-2. PMID: 38007464; PMCID: PMC10675866.

21. Karim SI, Irfan F, Saad H, Alqhtani M, Alsharhan A, Alzhrani A, Alhawas F, Alatawi S, Alassiri M, M A Ahmed A. Men’s knowledge, attitude, and barriers towards emergency contraception: A facility based cross-sectional study at King Saud University Medical City. PLoS One. 2021 Apr 26;16(4):e0249292. doi: 10.1371/journal.pone.0249292. PMID: 33901184; PMCID: PMC8075244.

22. Katatsky ME. The Health Belief Model as a conceptual framework for explaining contraceptive compliance. Health Ed Mono. 1977; 5(3):233–243

23. Hall, K. S. (2011). The Health Belief Model Can Guide Modern Contraceptive Behavior Research and Practice. Journal of Midwifery & Women’s Health, 57(1), 74–81. 10.1111/j.1542-2011.2011.00110.x

24. Lopez LM, Tolley EE, Grimes DA, Chen-Mok M. Theory-based strategies for improving contraceptive use: a systematic review. Contraception. 2009;79(6):411-417

25. Ritchie J, Spencer L. Analyzing qualitative data. Qualitative data analysis for applied policy research. Bryman A, Burgess RG (ed): Routledge, London. 1994:173-194.

26. Karim, S. I., Irfan, F., Saad, H., Alqhtani, M., Alsharhan, A., Alzhrani, A., … & MA Ahmed, A. (2021). Men’s knowledge, attitude, and barriers towards emergency contraception: A facility based cross-sectional study at King Saud University Medical City. PloS one, 16(4), e0249292

27. Boadu I. Coverage and determinants of modern contraceptive use in sub-Saharan Africa: further analysis of demographic and health surveys. Reprod Health. 2022;19(1):1–11. doi: 10.1186/s12978-022-01332-x

28. Wuni C, Turpin CA, Dassah ET. Determinants of contraceptive use and future contraceptive intentions of women attending child welfare clinics in urban Ghana. BMC Public Health. 2017;18(1):1–8.

29. Rominski SD, Morhe ES, Maya E, Manu A, Dalton VK. Comparing women’s contraceptive preferences with their choices in 5 urban family planning clinics in Ghana. Glob Heal Sci Pract. 2017;5(1):65–74. doi: 10.9745/GHSP-D-16-00281.

30. Albezrah NA. Use of modern family planning methods among Saudi women in Taif, KSA. Int J Reprod Contraception Obstet Gynecol. 2015;44:990–4.

31. Alomair N, Alageel S, Davies N, Bailey JV. Factors influencing sexual and reproductive health of Muslim women: a systematic review. Reprod Health. 2020;17(1):1–15.

32. Sedlander E, Yilma H, Emaway D, Rimal RN. If fear of infertility restricts contraception use, what do we know about this fear? An examination in rural Ethiopia. Reprod Health. 2022 Jun 13;19(Suppl 1):57. doi: 10.1186/s12978-021-01267-9. PMID: 35698228; PMCID: PMC9195198.

33. Sait, Moataz; Aljarbou, Abdullah; Almannie, Raed; Binsaleh, Saleh. Knowledge, attitudes, and perception patterns of contraception methods: Cross-sectional study among Saudi males. Urology Annals 13(3):p 243-253, Jul–Sep 2021. | DOI: 10.4103/UA.UA_42_20).

34. Al Riyami, A., Afifi, M., & Mabry, R. M. (2004). Women’s string-nameonomy, education and employment in Oman and their influence on contraceptive use. Reproductive health matters, 12(23), 144-154

35. Aneblom G, Larsson M, Odlind V, Tydén T. Knowledge, use and attitudes towards emergency contraceptive pills among Swedish women presenting for induced abortion. BJOG. 2002 Feb;109(2):155-60. doi: 10.1111/j.1471-0528.2002.01239.x. PMID: 11888097.]

36. Karim SI, Irfan F, Rowais NA, Zahrani BA, Qureshi R, Qadrah BH. Emergency contraception: Awareness, attitudes and barriers of Saudi Arabian Women. Pak J Med Sci. 2015 Nov-Dec;31(6):1500-5. doi: 10.12669/pjms.316.8127. PMID: 26870124; PMCID: PMC4744309.

37. Gupta N, Katende C, Bessinger R. Associations of mass media exposure with family planning attitudes and practices in Uganda. Studies in Family Planning. 2003;34(1):19–31. 10.1111/j.1728-4465.2003.00019.x.

38. Pazol K, Zapata LB, Tregear SJ, Mstring-nameone-Smith N, Gavin LE. Impact of Contraceptive Education on Contraceptive Knowledge and Decision Making: A Systematic Review. Am J Prev Med. 2015 Aug;49(2 Suppl 1):S46-56. doi: 10.1016/j.amepre.2015.03.031. PMID: 26190846; PMCID: PMC4532374.

39. Pazol K, Zapata LB, Dehlendorf C, Malcolm NM, Rosmarin RB, Frederiksen BN. Impact of Contraceptive Education on Knowledge and Decision Making: An Updated Systematic Review. Am J Prev Med. 2018 Nov;55(5):703-715. doi: 10.1016/j.amepre.2018.07.012. PMID: 30342633; PMCID: PMC10521032.

